# Trends and gaps in the use of citizen science derived data as input for species distribution models: a quantitative review

**DOI:** 10.1101/2020.06.01.127415

**Authors:** Mariano J. Feldman, Louis Imbeau, Philippe Marchand, Marc J. Mazerolle, Marcel Darveau, Nicole J. Fenton

## Abstract

Citizen science (CS) currently refers to some level of volunteer participation in any discipline of scientific research. Over the last two decades, nature-based CS has flourished due to innovative technology, novel devices, and widespread digital platforms used to collect and classify species occurrence data. For scientists, CS offers a low-cost approach of collecting species occurrence information at large spatial scales that otherwise would be prohibitively expensive. We examined the trends and gaps linked to the use of CS as a source of data for species distribution models (SDMs), in order to propose guidelines and highlight solutions. We conducted a quantitative literature review of 224 peer-reviewed articles to measure how the representation of different taxa, regions, and data types have changed in SDM publications since the 2010s. Our review shows that the number of papers using CS for SDMs has increased at approximately double the rate of the overall number of SDM papers. However, disparities in taxonomic and geographic coverage remain in studies using CS. Western Europe and North America were the regions with the most coverage (71.2%). Papers on birds (51.2%) and mammals (26.2%) outnumbered other taxa. Among invertebrates, flying insects including Lepidoptera and Odonata received the most attention. Compared to studies on animal taxa, papers on plants using CS data remain rare. Although the aims and scope of SDM papers are diverse, conservation remained the central theme of SDM using CS data. We present examples of the use of CS and highlight recommendations to motivate further research, such as combining multiple data sources and promoting local and traditional knowledge. We hope our findings will strengthen citizen-researchers partnerships to better inform SDMs, especially for less-studied taxa and regions. Researchers stand to benefit from the large quantity of data available from CS sources to improve global predictions of species distributions.

## Introduction

Species distribution models have become a widely used tool in ecology in recent years [1-3]. Understanding the association between the occurrence of species and environmental conditions is a first step in addressing questions about species distributions, abundances and habitat preferences [4, 5]. Current global-scale issues such as climate and land-use changes have increased the need to understand and predict the distribution of migratory or invasive species across a landscape. In fact, knowledge on species distribution is paramount to develop biodiversity conservation and management strategies [6]. The fundamental theory behind species distribution models (SDMs, hereafter) assumes that the presence of a species in a given location strongly depends on the environment, which implies that ecologists are able to estimate future species distributions based on the environment of current locations [7]. Specifically, SDMs use empirical data to link information about the presence of a species to the environmental variables of their known locations, and apply statistical models to predict the spatial distribution of species [4, 5, 8]. Consequently, we can identify three major components in any framework for SDMs: species presence data, landscape or environmental data, and a statistical model that links the first two components.

Species distribution models have been used to tackle a wide range of scientific issues at different spatial and temporal scales. SDMs are used in both fundamental science and applied sciences in biogeography, evolution, dispersal, migration, species invasion, meta-population, conservation, and climate change [3]. For example, SDMs have shown their value, for example, in characterizing the current distribution range of a species [9, 10], predicting variables measured in the field [11], assessing species invasions [12], or evaluating the impact of land-use changes [13]. Species distribution models have multiple uses, including predicting spatial changes in response to climate change [14, 15], assessing the suitability of possible conservation areas [16, 17], or suggesting areas to improve survey efforts for rare species [18].

Modelling the spatiotemporal distribution of species usually requires a large amount of information collected over multiple years of standardized fieldwork. However, the long-term collection of broad-scale information on a wide range of species is prohibitively expensive. Yet, for some taxa, an impressive volume of data collected using non-standardized protocols is currently available in museum collections, distribution atlases and online portals, through efforts collectively labelled as citizen science (CS, hereafter). Globally, a huge variety of CS programs are currently being implemented involving a wide range of taxa [19]. Nevertheless, CS data are still challenging to analyze due to the intrinsic issues of non-standardized protocols that can affect the credibility and quality of the data.

Issues within CS datasets arise from the large number of observers that collect species data. Previous studies have tackled the different sources of variation in CS data [20-22]. Firstly, CS datasets are typically biased towards human population centers, areas that are easy to access, protected areas, or regions frequented by active observers. These problems lead to disparities in effort between over-sampled and under-sampled areas [22-27]. Secondly, geographical coverage of CS data can be biased towards well-financed and more industrialized countries, mainly in North America and Europe [28-30]. These two regions contribute substantially more data than any other region in the Global Biodiversity Information Facility (GBIF) database [31-35]. Consequently, a large proportion of samples occur in a restricted geographical extent, controlled by administrative borders. This results in a non-representative sample of species’ distribution. Thirdly, over time, the observation and reporting protocols can change. For example, the Audubon Christmas Bird Count at its start in 1900 aimed at offering an alternative to hunting on Christmas Day morning, with a loose survey protocol. The count day became flexible over years. For example, it was from December 22-29 in 1940-41 [36] and December 21 to January 2 in 1966-67 [37]. In 1966-67, the goal became to collect a snapshot of wintering birds around Christmas time: the survey protocol was standardized [38]. In 2000, the survey period expanded again, this time to 23 days as the count should now be completed on a day between December 14 and January 5 [39]. Unfortunately, changes in survey protocols are often poorly documented. Fourthly, among biological groups, CS observations can be taxonomically biased because volunteers are usually attracted to large and common species, to species that are brightly colored and easy to detect, and to more charismatic groups [21, 34, 40, 41]. This taxonomical disparity results in more information on relatively well-known groups than for under-reported groups. Finally, another source of variation in CS programs includes the variation in skill and expertise among observers, primarily due to the participation of a wide range of volunteers. As the quality of observations depends on the ability of observers to correctly identify species, this introduces a qualitative bias that can lead to misinterpretation of results. Indeed, this inter-observer sampling variation increases for species that are harder to detect [42-44]. Bias inherent to each of these five sources of variation may influence predictions of future trends. A major challenge is to account for these issues in species distribution models [45, 46].

Despite these issues regarding data quality, the use of CS has increased in recent years in different fields of study [47, 48]. For instance, CS is used in astronomy to classify galaxy images or to search for signals in radio data, and in atmospheric sciences to record the quality of air, soil, and water [47, 49, 50]. However, the main application of CS is in conservation and ecology to monitor species occurrence [33, 51]. Several reviews have focused on how CS contributes to biodiversity monitoring [33], global change [52], and conservation biology [53]. Considering the increasing prevalence of CS in ecological studies, there is a need to synthetize the application of CS in SDMs. Accordingly, the main objective of this review was to quantify the variation over time in the use of CS data as an input for modelling species distribution. To achieve this objective, we assessed the current strengths in the use of CS in SDMs and identified partiality and gaps relative to taxa, regions, and data acquisition methods. Our main questions were: (1) What is the trend in the use of CS data for SDMs over the last decade? (2) Is there variation across regions, taxa, and types of data used? (3) What are the information gaps and how can we meet research needs in the near future?

Given the increasing use of citizen science in different field studies, we expected an increasing use of CS in SDMs. We also anticipated that because volunteers behave differently according to the region and group of interest, the set of papers would reflect clear preferences towards regions that are easy to access and groups that are apparently visually appealing to volunteers. However, we expected that these preferences would change over time due to the growing diversification of initiatives and platforms worldwide over the last decade.

## Materials and methods

### Paper selection

We used the Scopus search engine to conduct a literature review of peer-reviewed papers focusing on species distribution models that used citizen science. Our search spanned a period of 10 years, considering papers from 2010, when the “citizen science” term was widely accepted by several authors [50, 54, 55], until 17 October 2019. We searched for papers using the following combination of keywords: (“citizen science” OR “public participation” OR “community monitoring program” OR “participatory monitoring”) AND (“species distribution model” OR “predictive model” OR “distribution map” OR “invasive species” OR “occupancy model” OR “occurrence” OR “migration” OR “climate change”).

### Data collection

From the first Scopus search, we screened a total of 3,836 papers based on their title, abstract and keywords (Fig 1a). We dismissed those not related with either CS or SDMs. The remaining 800 papers were reduced after a further revision of abstracts and methodologies. We excluded papers written in languages other than English (n=4), all review papers, and also papers using data gathered by volunteers but without applications of SDMs (e.g., first report of a species or new occurrence data). To consider a given paper as relevant for our review, each of the following three conditions had to be met: the data included the presence or abundance of a biological group, the data were collected by volunteers (either partially or entirely), and a statistical method was applied to assess relationships with environmental data (Fig 1a). We ended up with 224 papers in peer-reviewed journals that formed the basis of the analyses presented herein. Details and extracted information about all papers included in our review are listed in Supporting information (S1 Table). We believe this list does not reflect the total influence of CS programs in SDMs, but only their contribution to published articles.

**Fig 1.**
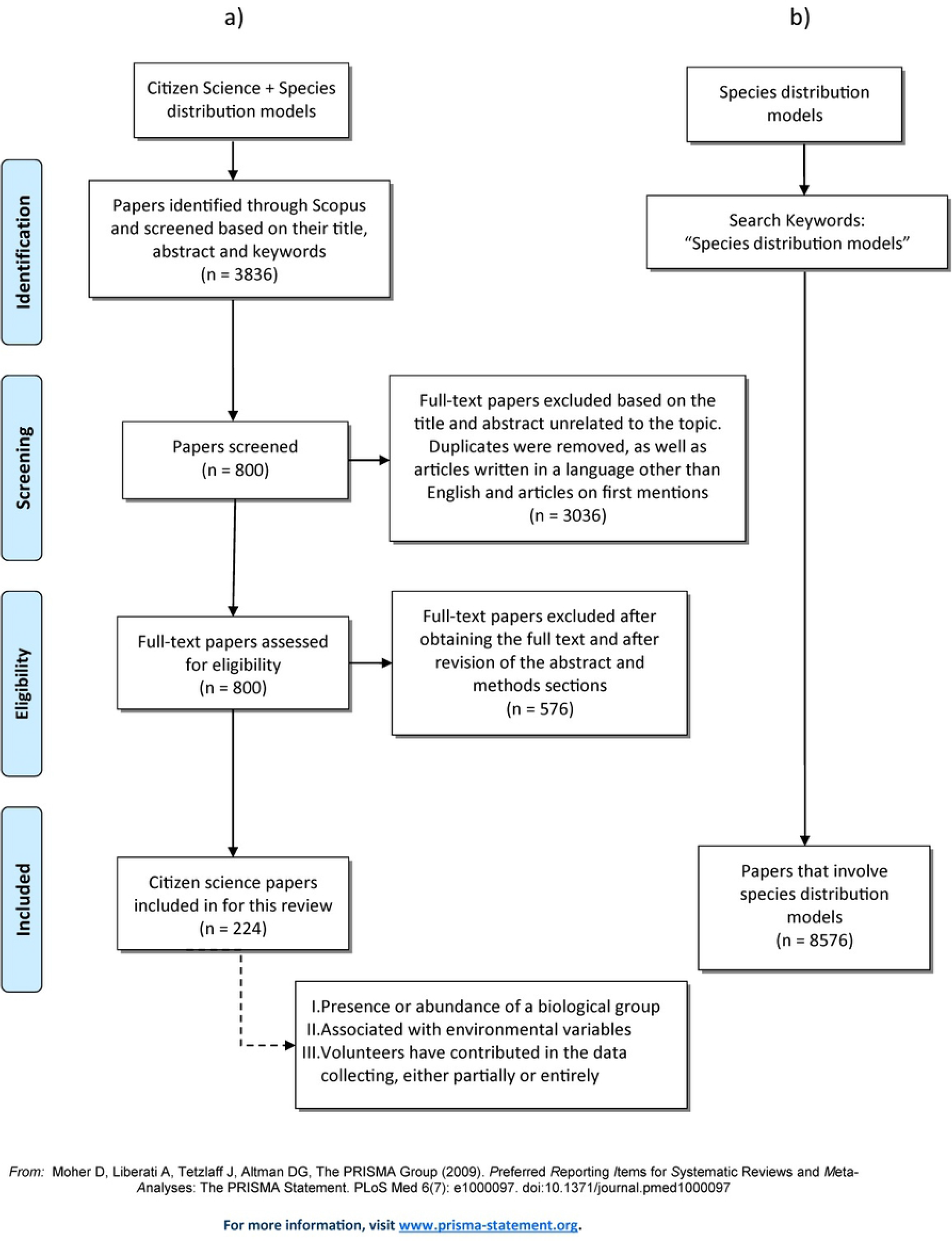
Flow chart of papers selection for a) the CS papers set and b) for the entire SDMs field set. All 224 papers in a) are listed in S1 Table.

From each paper we extracted the following information: (1) year of publication; (2) focal taxa; (3) source of data or platform used (if any); (4) country and region where the data was taken; (5) scope of the SDM paper — when appearing in the title, abstract, or keywords; (6) data type used (presence-only, presence-absence, or abundance); (7) statistical approach used, and 8) the method of collecting CS data (opportunistic data, count data, community-based monitoring, historical records, local knowledge, or trained volunteers). In order to assess the contribution of CS to SDMs over the last decade, we first compiled papers within Scopus by using the keyword “species distribution models” to obtain the number of papers in this field (Fig 1b). This search resulted in 8576 papers.

### Data Analyses

#### Contribution of CS to the SDMs

We consider these papers to be the number of papers published within the SDM field in the last decade. We tested for differences in the number of CS-SDM papers and the overall number of SDM papers using linear models (with the number of papers on a log scale) that included an interaction term between the year and type of paper (CS-SDM and SDMs). We expected papers using CS data to have increased at a faster rate than the SDM field as a whole.

#### Taxonomic groups

In order to assess the representation of CS within the biological groups, we categorized each paper within the following taxonomic groups: invertebrates, plants and fungi (including bryophytes and lichens), fish, reptiles, amphibians, mammals, and birds. We used a chi-square test to compare the number of CS papers with data on each group observed to the numbers expected based on the proportion of species in each group according to the Catalogue of Life [56] (accession date April 2020). To compare the observed (CS) and expected proportions for each of the five taxonomic groups, we constructed a logistic regression model excluding the intercept to estimate the logit of the probability that a taxa *t* appears in a CS study:

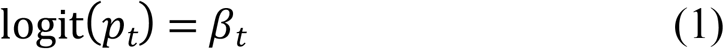

We then calculated a Z-score from the difference between β_t_ and the logit of E_t_, the expected proportion for that taxa, scaled by the standard error of β_t_:

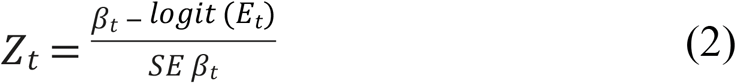

We obtained a two-tailed p-value for the null hypothesis that the observed proportion was equal to the expected proportion by comparing *Z_t_* to the standard normal distribution. We excluded papers that focused on more than one taxonomic group to meet assumptions of statistical independence of observations. For invertebrates, we only analyzed the taxonomic orders represented in our CS set of papers (Lepidoptera, Odonata, Hymenoptera, Coleoptera and Mollusca).

#### Geographic regions

Papers were individually classified into country and continent of origin of the CS data, including Africa, Asia, Eastern Europe, Western Europe, Oceania, North America, Central America, and South America. To assess if these regions were over or under-sampled in the CS papers set, we used a one-sample chi-square test to compare the number of CS papers in each region to the number expected based on the proportion of the Earth’s land area covered by each region (from http://www.worlddata.info; accessed 29.11.19). Then, we compared these observed and expected proportions for each of these geographic regions. Using the same strategy as above, we constructed a logistic regression model that excluded the intercept to estimate the logit of the probability that the region _*y*_ appears in a CS study:

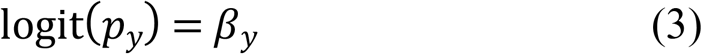

We then calculated a Z-score from the difference between β_*y*_ and the logit of E_*y*_, the expected proportion for that region, scaled by the standard error of β_y_:

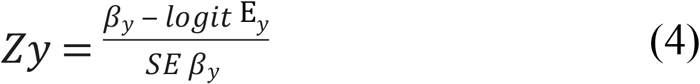

We compared the Z_*y*_ against the standard normal distribution. We excluded papers that focused on more than one region to meet assumptions of statistical independence of the observations.

## Results and discussion

### Year of publication

Our analysis indicates that the use of CS data in the peer-reviewed SDM literature has increased in frequency over the past 10 years (Fig 2a). Numerous authors have indicated the increase in publications using different types of CS data [47, 48, 52, 57, 58], but also the growing rate of SDMs in publications [59, 60]. In our analysis, however, the use of CS in SDMs is growing approximately twice as fast as the number of papers using SDMs in general (Fig 2b). In addition, given its peak in 2019 with 75 papers (Fig 2a), the next few years may extend the exponential growth of the use of CS in SDMs.

**Fig 2.**
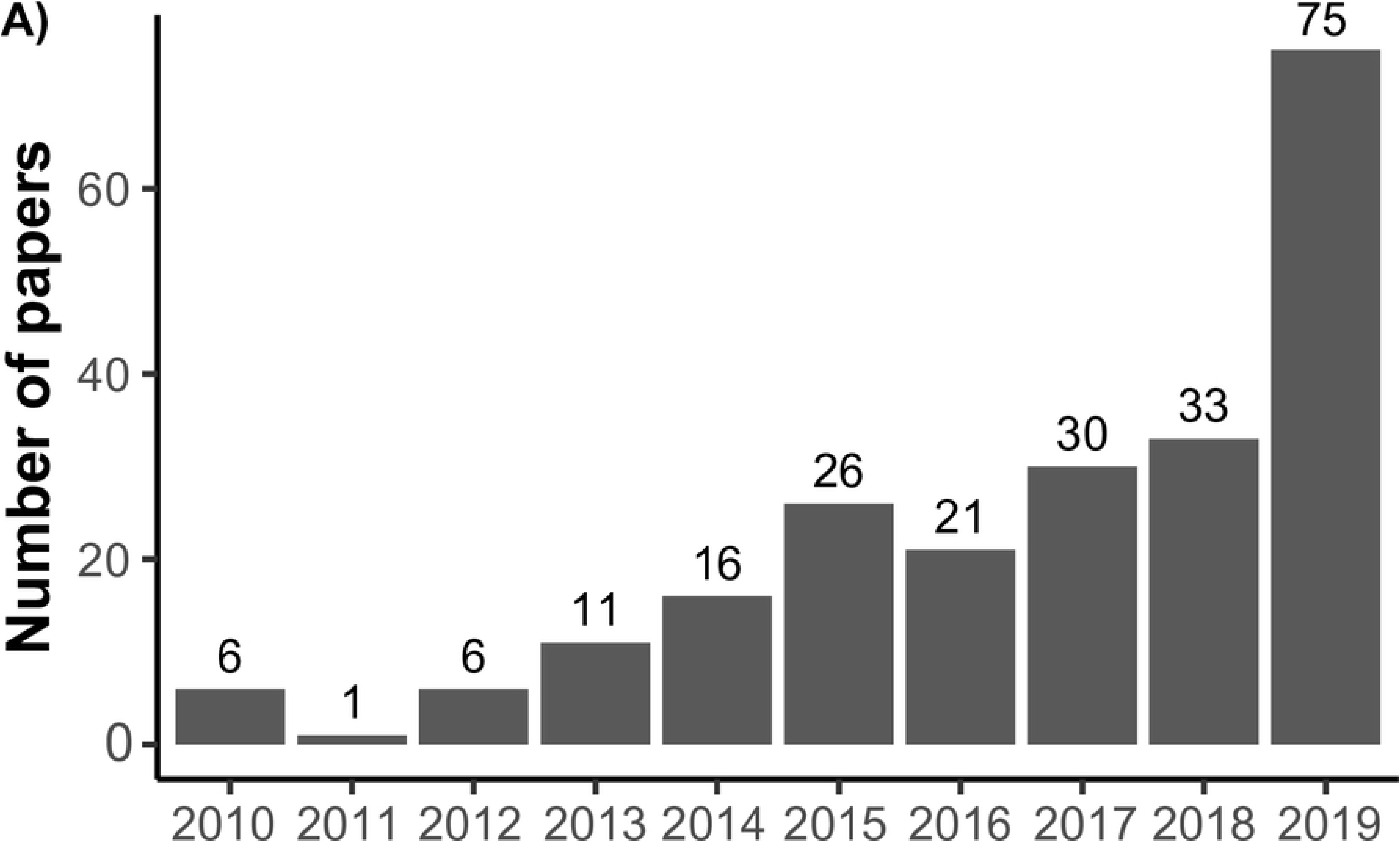
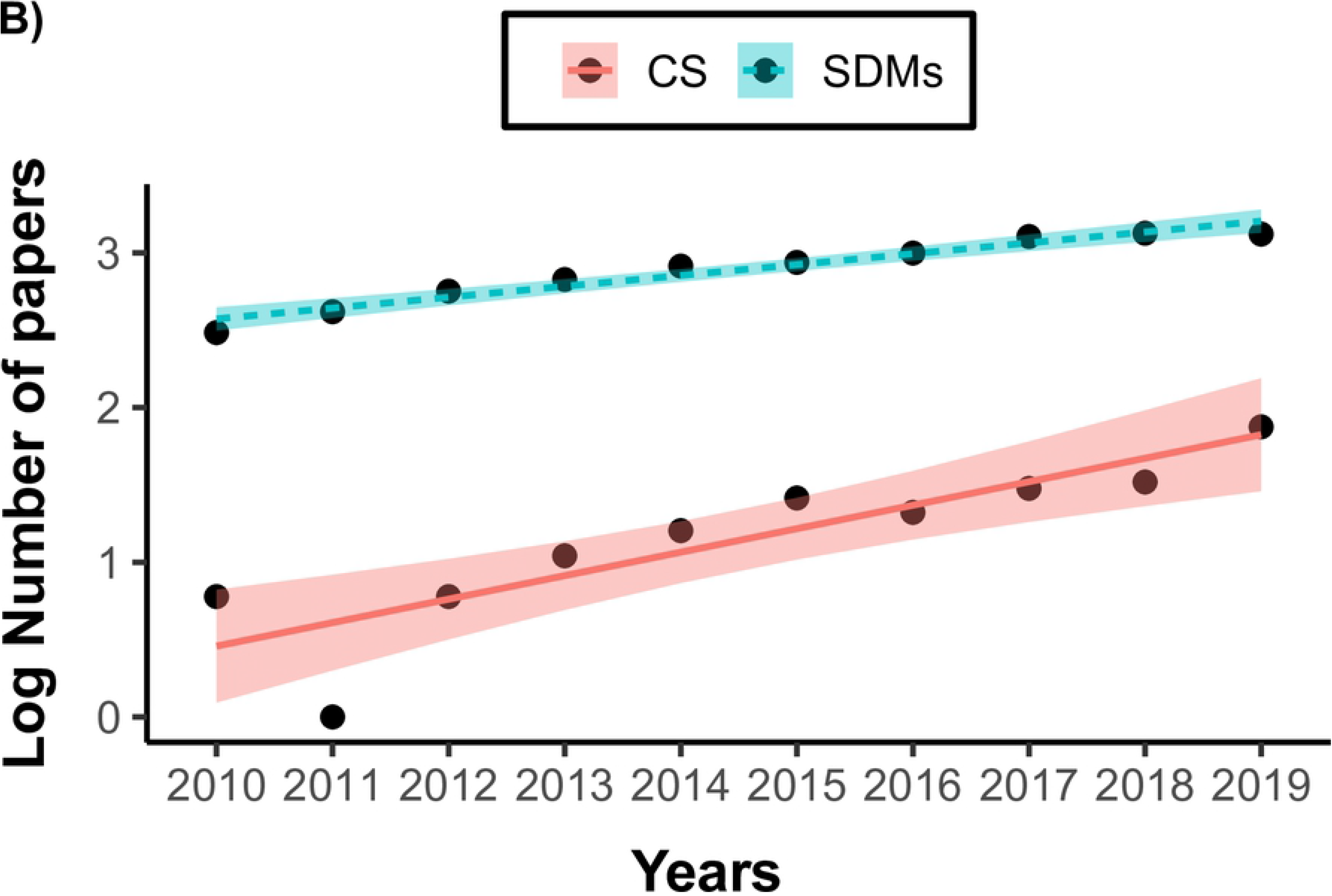
(a) Annual number of papers that have used species distribution models (SDMs) with citizen science (CS) data; (b) Linear regression of the log of total papers using SDMs (blue) and the papers using CS data (red) across the 10-year period covered by our review (difference in slopes: −0.08, std. error: 0.03 p=0.01).

### Taxonomic groups

As we predicted, there were marked variations among taxonomic groups, with birds (n = 87), invertebrates (n = 49), and mammals (n = 43) being the main taxa studied. Reptiles (n = 12), fish (n = 11) and amphibians (n = 11) received less attention (Fig 3a). This taxonomic preference towards bird species was previously noted by other authors [33, 47, 52].

**Fig 3.**
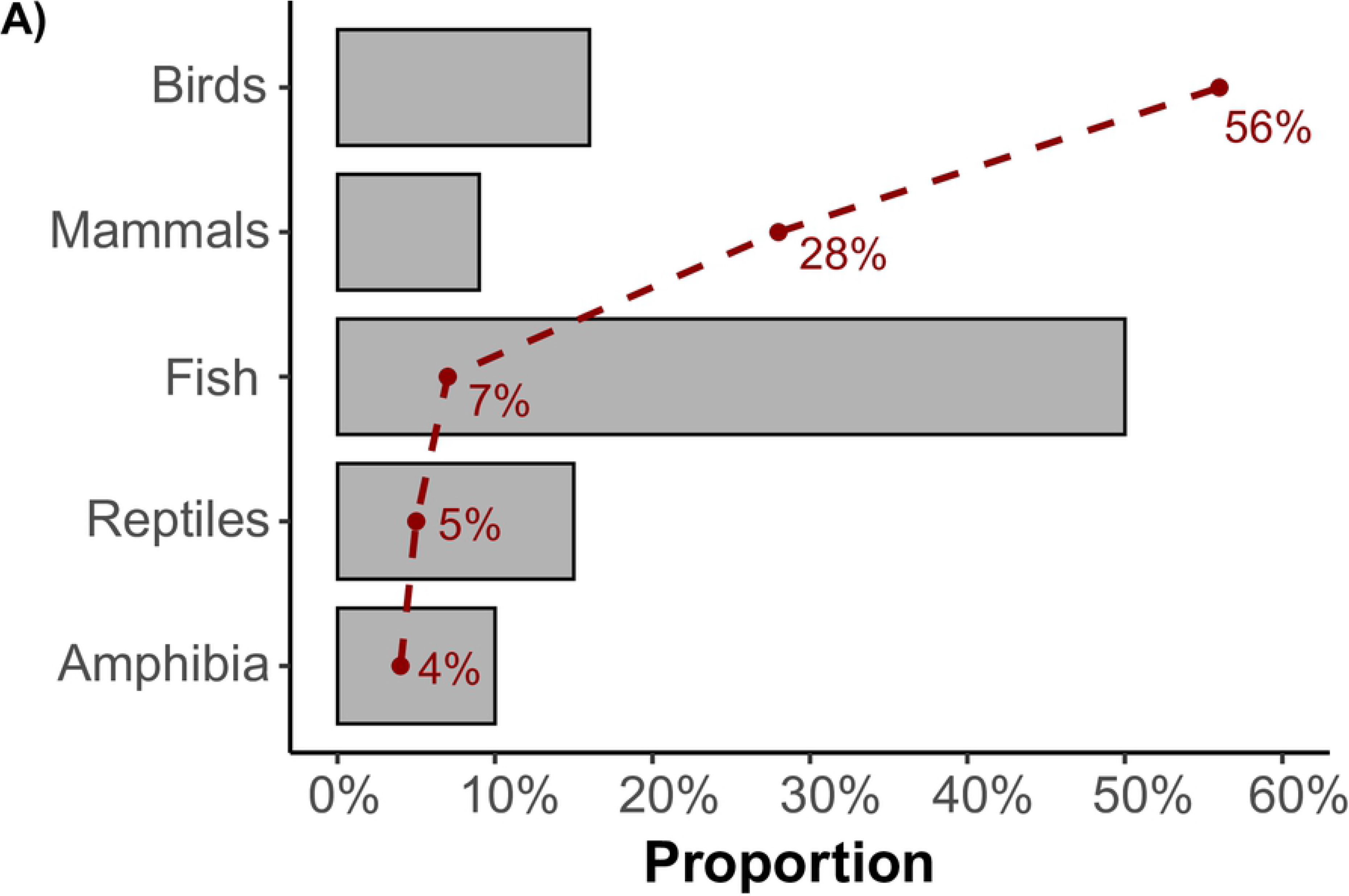
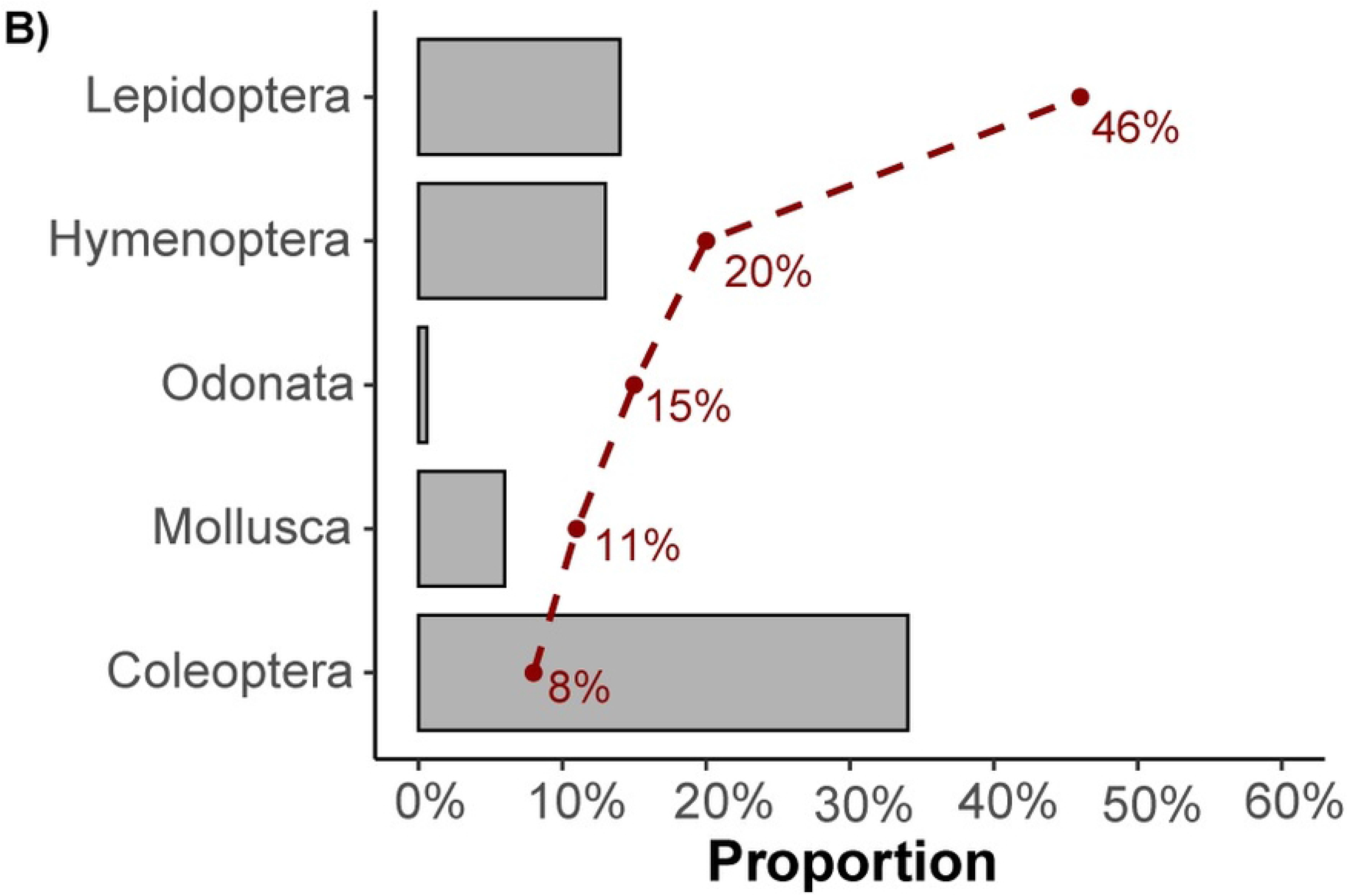
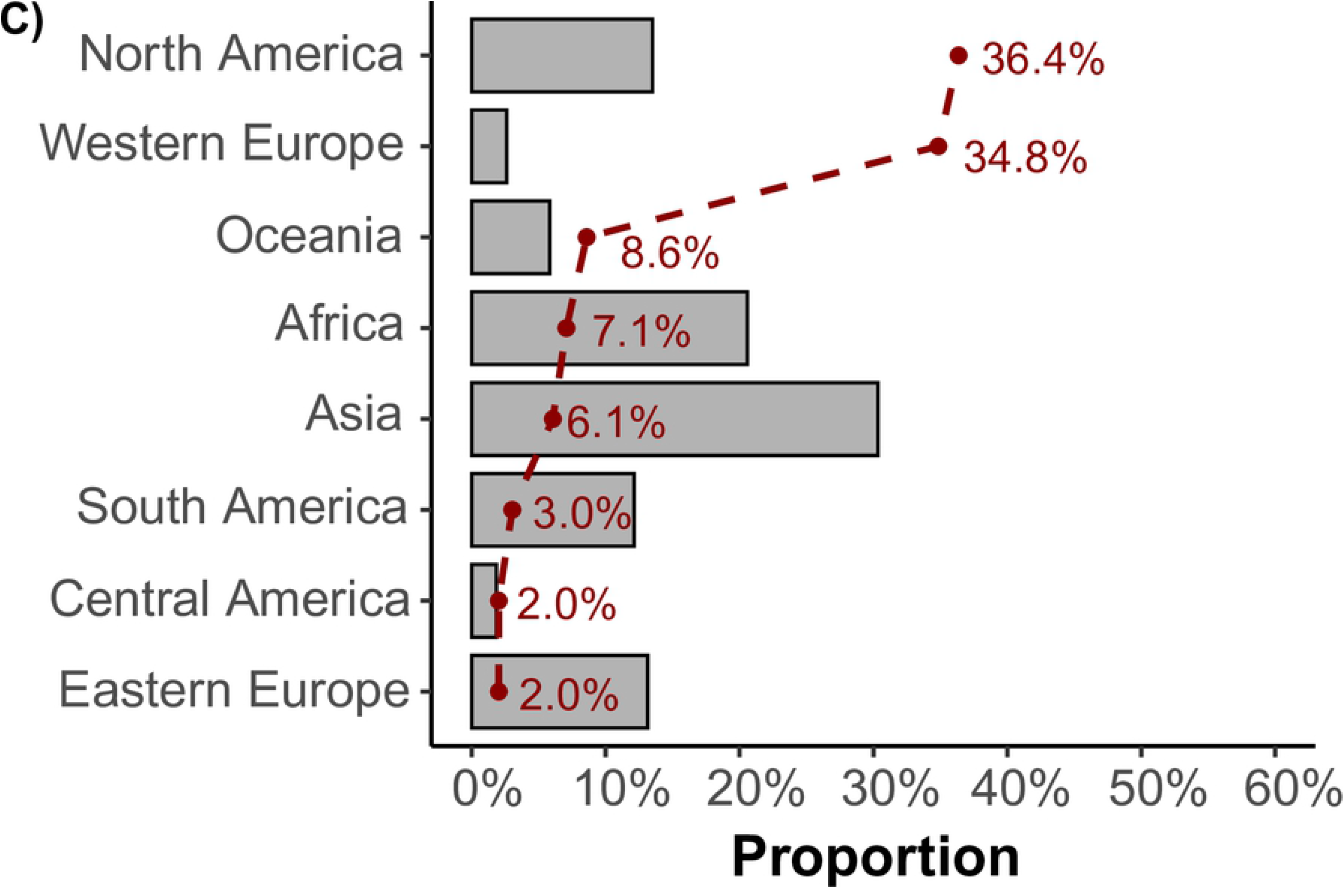
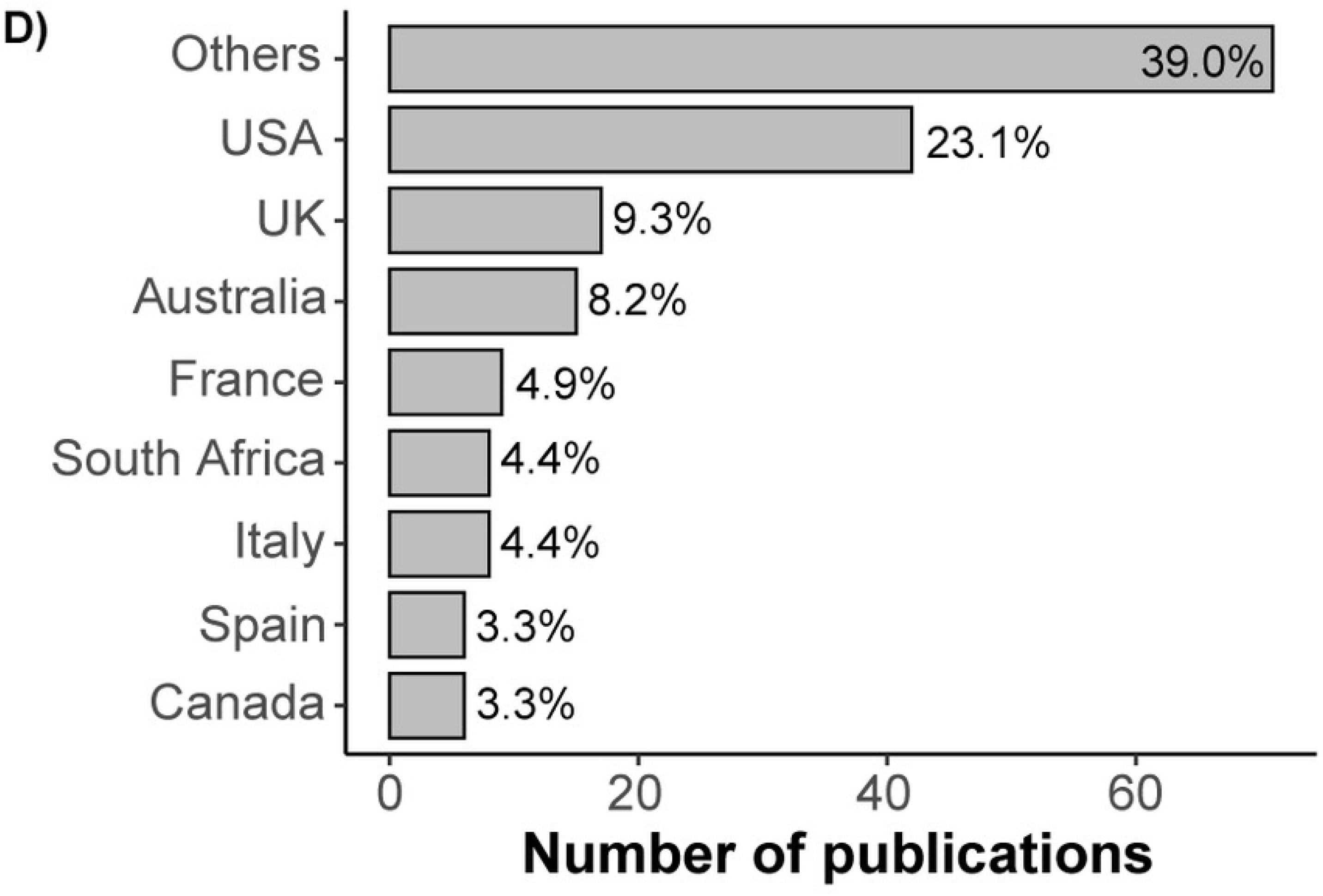
Proportion of citizen science (CS) papers from this review (red dotted line) relative to the proportion of global richness in the Catalogue of Life (grey bars) by taxa, for (a) Vertebrates (n = 151) and (b) Invertebrates (n = 41); (c) Proportion of CS papers by data collection region (red dotted lines) relative to each region’s fraction of the Earth’s land area (grey bars); and (d) Proportion of CS papers by country.

Compared to their global richness, vertebrate groups were unequally represented in the CS papers (χ^2^ = 280.14, df = 4, p < 0.05). Specifically, birds (*Z_birds_* = 11.8; p < 0.05) and mammals (*Z_mammals_* = 7.3; p < 0.05) were over-sampled, whereas amphibians (*Z_amphibians_* = −2.3; p < 0.05), reptiles (*Z_reptiles_* = −8.9; p < 0.05) and fish (*Z_fish_* = −8; p < 0.05) were under-represented in CS papers compared to their estimated global richness in the Catalogue of Life database (Fig 3a)Fig 3. We also found differences between the observed proportion of invertebrates in our CS data set and the proportion expected based on global richness (χ^2^ = 326.9, df = 11, p < 0.005). Lepidoptera (*Z_lepidoptera_* = 3.5; p < 0.05) and Odonata (*Z_odonata_* = 5.8; p < 0.05) were over-sampled relative to their global richness (Fig 3b). Only Coleoptera (*Z_coieoptera_* = −4.2; p < 0.05) were under-sampled. However, the proportion of papers on Mollusca (*Z_mollusca_* = 0.9; p = 0.39) and Hymenoptera (*Z_hymenoptera_* = 0.7; p = 0.44) did not differ from the proportion expected from global species richness. The remaining invertebrates orders in the Catalogue of Life database did not occur in the set of CS papers studied (Fig 3b).

The plant and fungi group included papers involving vascular plants (n = 18), fungi (n = 3), lichens (n = 1), and bryophytes (n = 1; S1 Table). Considering the known number of species in each group according to the Catalogue of Life database (vascular plants: 348,000 species; fungus: 140,000 species; bryophytes 16,000 species), plant taxonomic groups were remarkably under-represented in CS papers. The major obstacle could be that identifying plants up to species level in the field is sometimes complex, even for expert botanists [61, 62]. Plant identification is time consuming for several families, requires significant botanical skills, and can be frustrating for non-expert volunteers [62]. In addition, there is not as strong a tradition for botanists in sharing observations using online portals, compared to animal databases. Nonetheless, plant initiatives seem to be highly attractive to the general public. Millions of observations are produced and stored in broad databases such as GBIF, iNaturalist, and in particular botanic platforms such as Pl@ntNet [63], Project Bud Burst [64], or Plant Watch Canada [65]. Several authors recommend using this information collected from volunteers for the early detection and control of invasive species [66, 67], or to improve the performance of models [67-69]. Nevertheless, our review confirms a notable under-use of plant, fungi, lichen and bryophyte public databases in the last decade in papers that model the distribution of species (10% of the total).

### Geographic coverage

Our review identified strong geographic biases in CS sampling efforts (χ^2^ = 1424.5, df = 7, p < 0.05). While Western Europe (*Z_Western Europe_* = 21; p < 0.05) and North America (*Z_North America_* = 8.1; p < 0.05) were over-sampled relative to their fraction of the planet’s land area, Africa (*Z_Africa_* = −4.5; p < 0.05), Asia (*Z_Asia_* = −6.3; p < 0.05), South America (*Z_South America_*= −3.6; p < 0.05) and Eastern Europe (*Z_Eastern Europe_* = −3.9; p < 0.05) were under-sampled (Fig 3c). Oceania (*Z_Oceania_* = 1.5; p =0.11) and Central America (*Z_Central America_* = 0.1; p = 0.89) were sampled proportionally to their area. At the country level, most of the papers using CS data were from USA (n = 42), the UK (n = 16), Australia (n = 15), France (n = 9), and South Africa (n = 8; Fig 3d).

Such a strong geographic inclination toward Europe and North America has already been indicated by several authors [33, 35, 70]. Others also revealed the same pattern of CS being predominantly conducted in Europe and North America, but with a greater number of studies in South and Central America [17, 30] than reported in our study. This large disparity of CS-based papers is likely influenced by three factors. First, North America and Europe host more developed countries, which have more funding available for research [40], and consequently tend to publish more. Some of these countries have traditional national platforms such as the National Biodiversity Gateway (NBN) in the United Kingdom (containing around 127 million records), the Atlas of Living Australia (ALA; containing 87,179,824 records on 19^th^ April 2020), or the Sweden Species Gateway (containing around 60,000 species). Individual country platforms share characteristics associated with successful CS programs that contributed more to global biodiversity monitoring. These platforms receive important support by national governments and are linked to well-funded institutions with active involvement of academic researchers [33]. These factors explain why the expansion of CS platforms in developing countries might be limited by the availability of necessary infrastructures [33]. Secondly, this geographic pattern is consistent with the tradition of CS, which emerged in North America and then spread globally, primarily driven by some iconic platforms and surveys such as the Christmas Bird Count, *eBird*, and Project BudBurst [64]. Lastly, in regions with fewer papers using CS data, sharing of biodiversity data remains difficult due to a lack of an open-access culture and language barriers [33, 71]. Papers in languages other than English were not included in our review, which may introduce a bias in our coverage of geographical areas in Arab countries, Latin America, or Asia [70]. In addition to language barriers and geographic location, national security concerns and economy also fosters spatial variations in the coverage of global databases [70].

### Source of data

The main reason for the predominance of bird CS papers was the significant use of three global networks of birders: the *eBird* project, the Breeding Bird Survey (BBS), and the Southern African Bird Atlas Project (SABAP, Fig 4). For insects, the Butterfly Monitoring Scheme (BMS) was widely used and proved to be a powerful tool to detect population trends [72]. Even if GBIF was the second most used source of information, this portal aggregates global biodiversity information from a variety of sources [33], including other CS portals listed in Fig 4. Indeed, the major GBIF contributor is *eBird* [33, 70]. For that reason, we cannot dissociate the GBIF database from other sources of CS in Fig 4.

**Fig 4.**
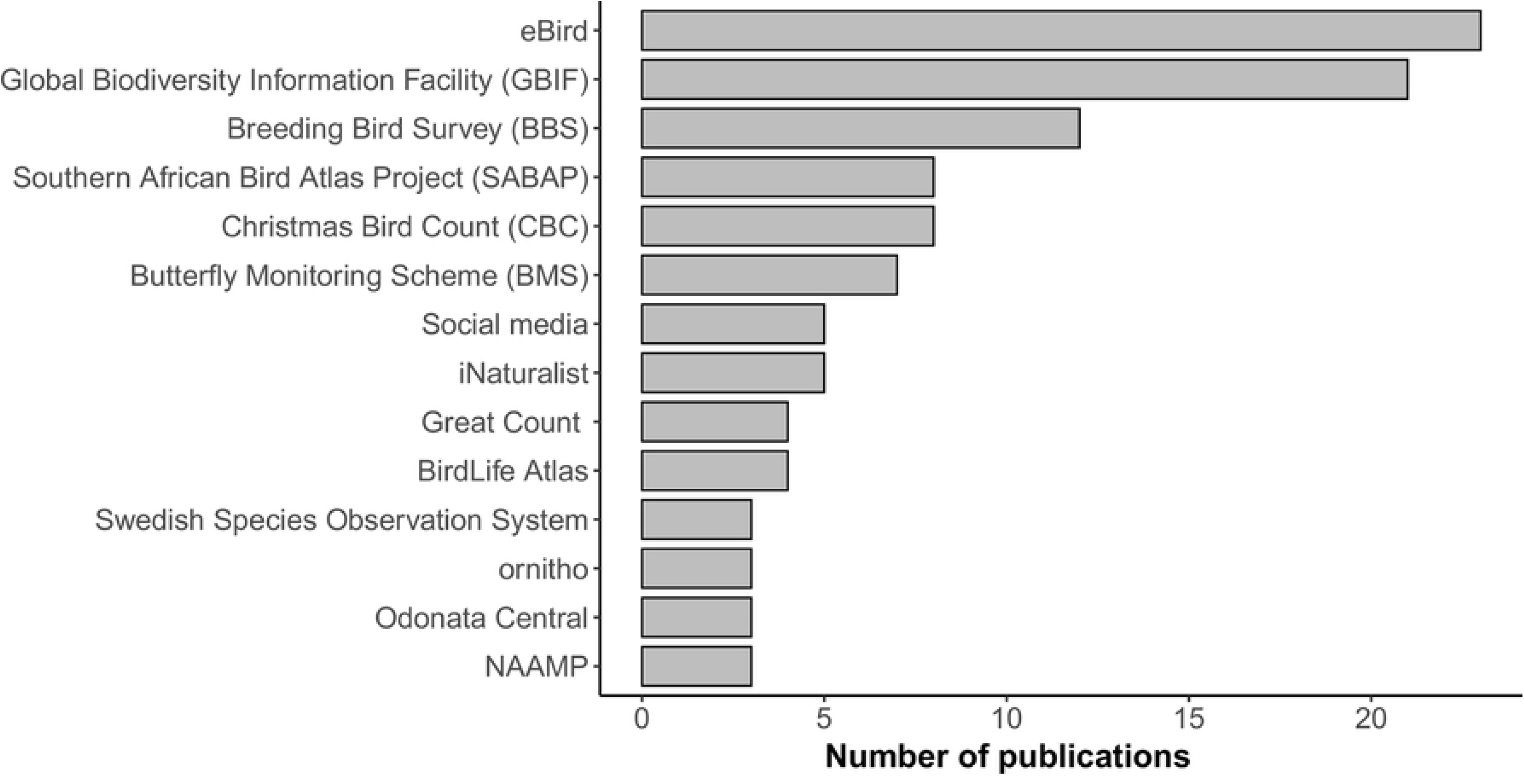
Sources of information used in the papers included in the literature review (n = 224). Only databases with three or more papers are shown. Great Count: world-wide surveys targeting birds and mammals.

### Study scope

Although the scope and geographical coverage varied greatly among SDM papers using CS data (Fig 5a), most papers addressed issues related to conservation (n = 104), followed by population trends (n = 69), habitat suitability (n = 61), and climate change (n = 57; Fig 5a). Furthermore, conservation remains the central aim of these studies (Fig 5b). The same pattern was observed for SDMs in tropical regions [1]. Most conservation papers using CS documented species of conservation concern, rare species, or poorly-studied regions [25, 73].

**Fig 5.**
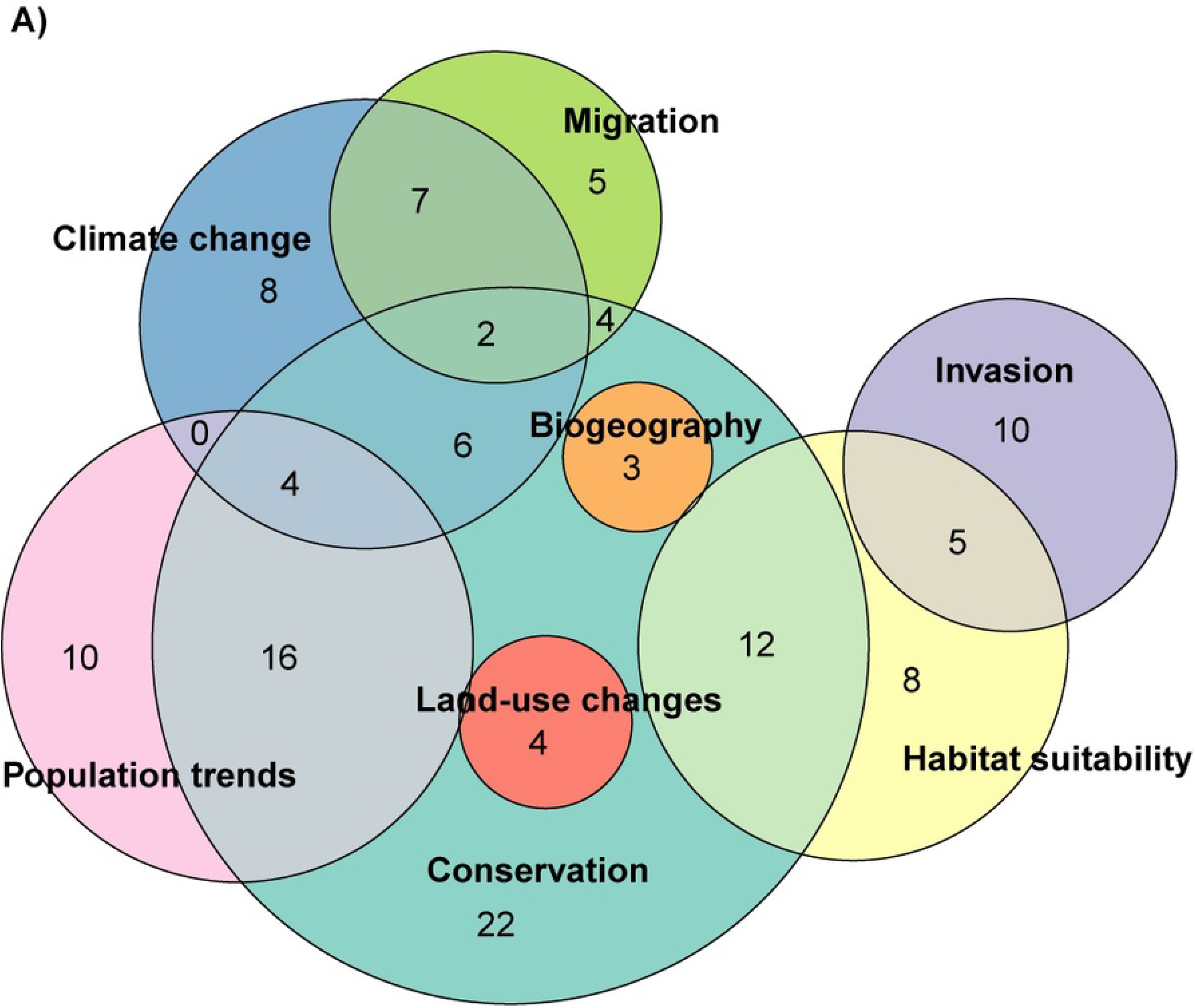
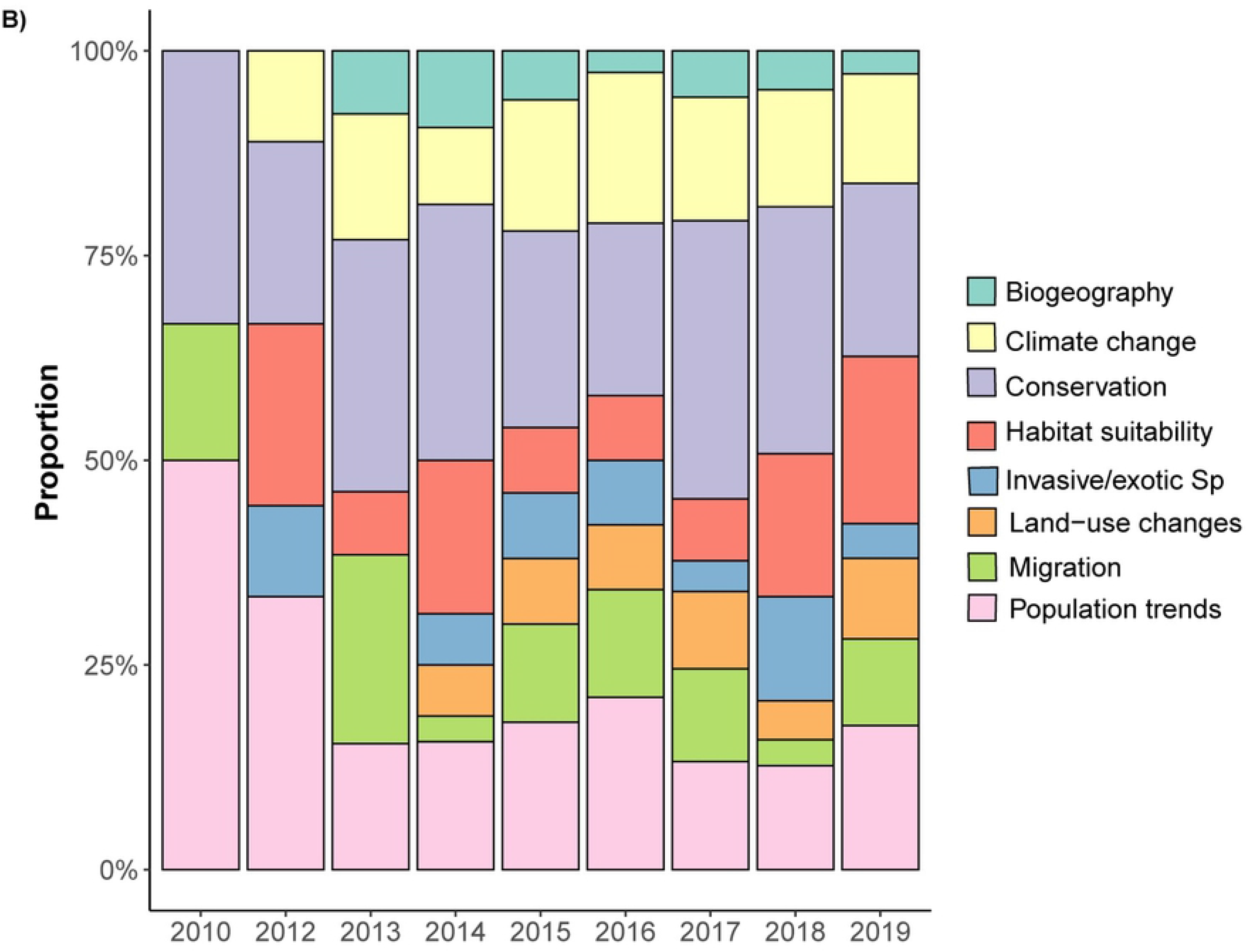
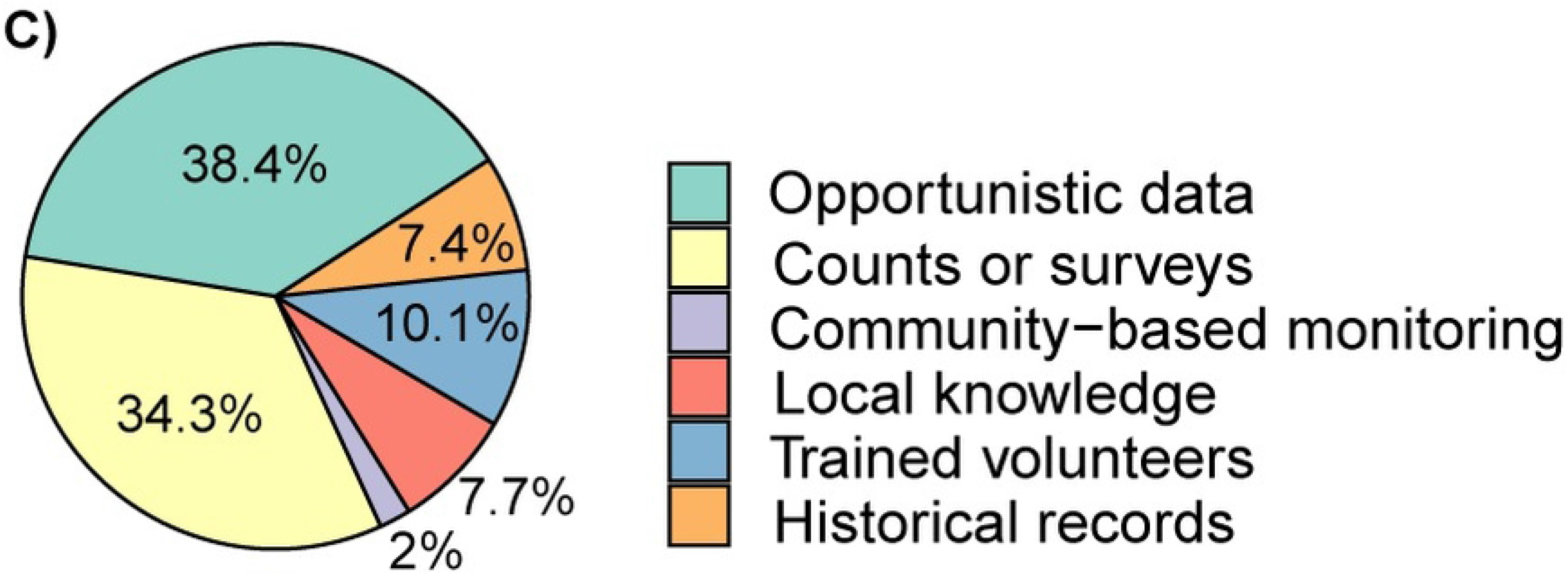
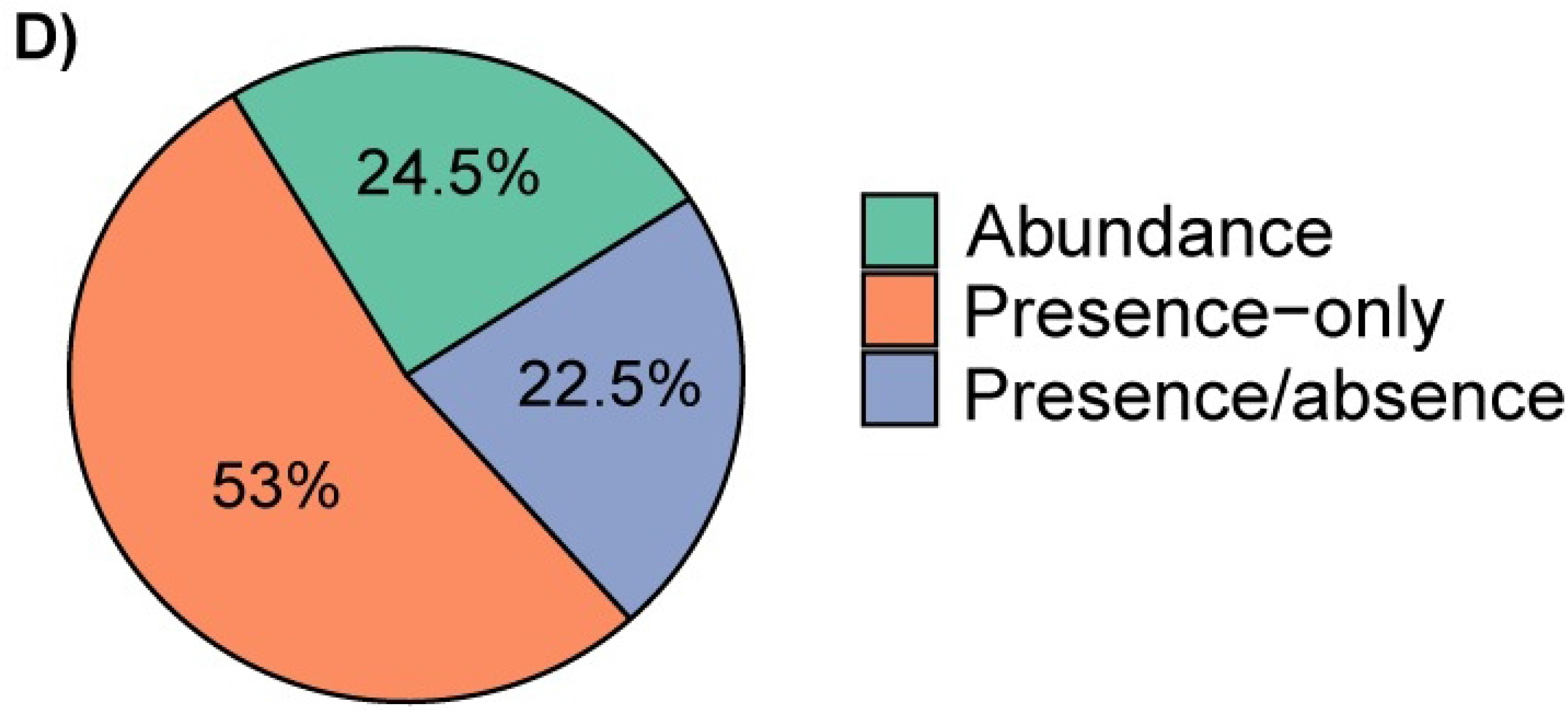
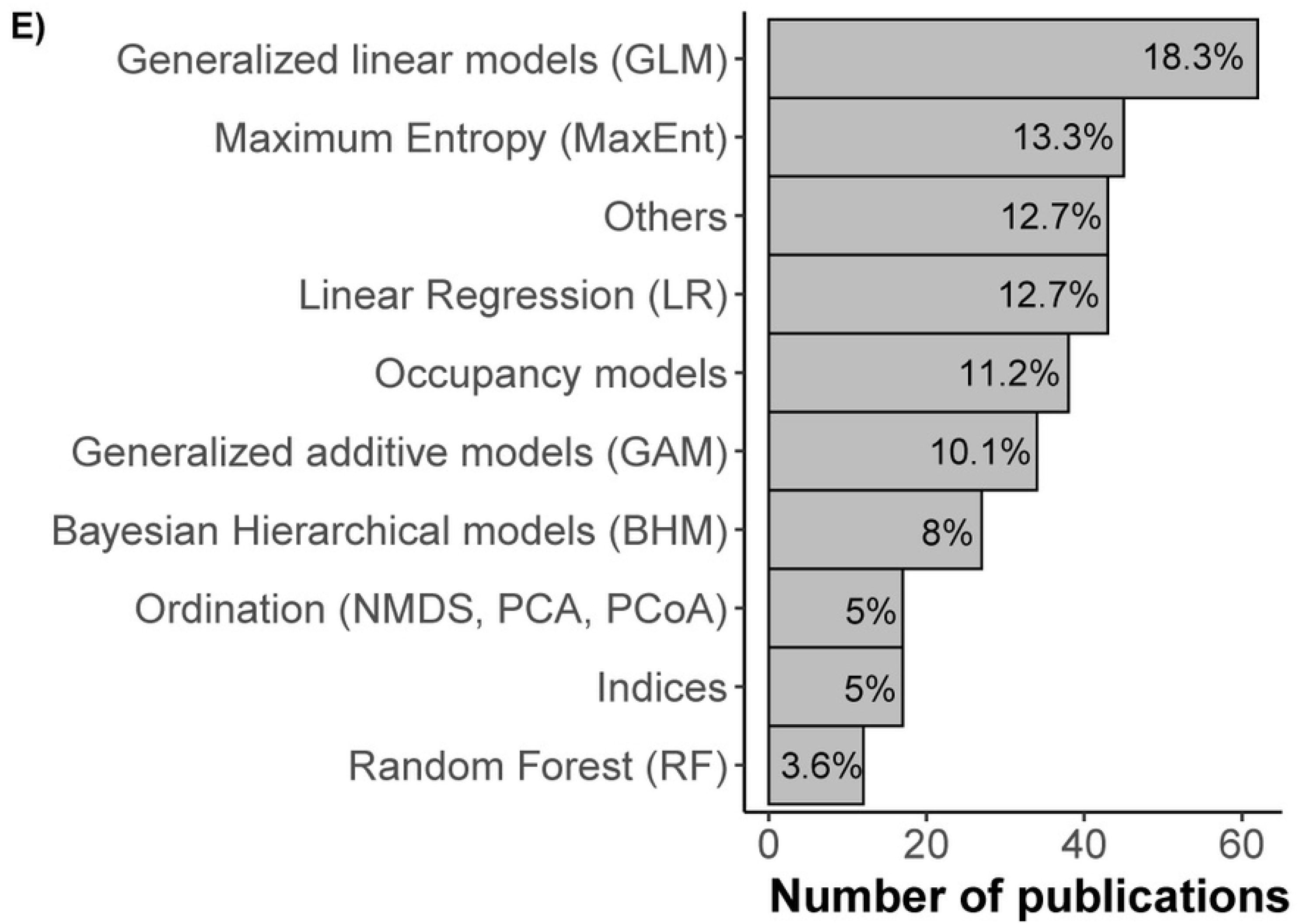
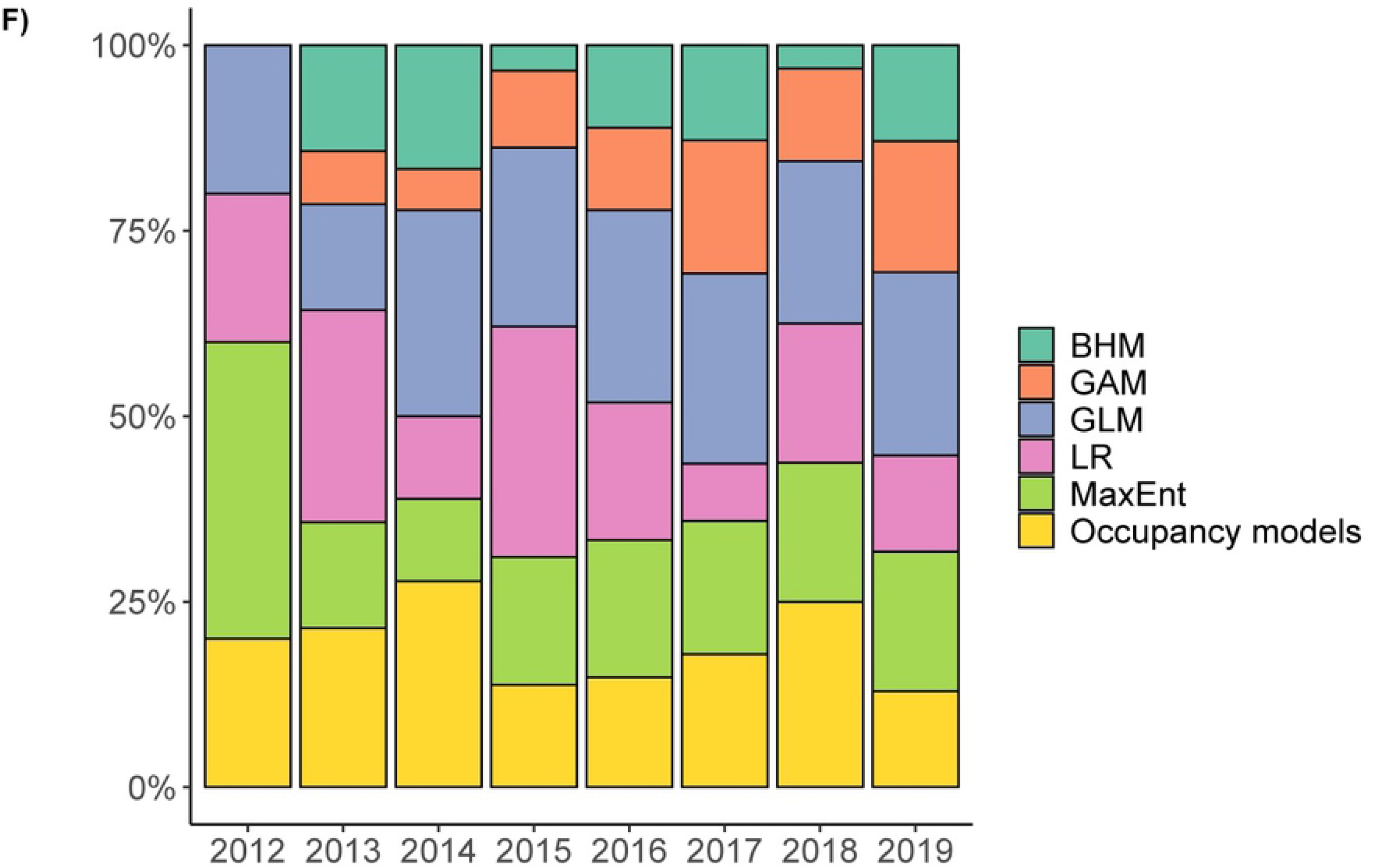
Papers that have analyzed species distribution models (SDMs) using citizen science (CS) data in our literature review illustrating (a) study scope illustrated by a Venn diagram (intersections containing a single study not shown); (b) study scope per year; (c) method of collecting citizen science data; (d) data type used; (e) statistical approach and (f) statistical approach per year. All details for the 224 papers are cited in Supplementary Information S1 Table.

### Method of collecting CS data

A number of methods of collecting CS data were reported in the set of papers we reviewed. Opportunistic data collection accounted for 38.4% of the papers (Fig 5c). With the growing popularity of online databases compiling occurrence data, the predominance of opportunistic data collection is not surprising [74]. Opportunistic data can be collected in many forms, including crowdsourcing databases or historical records from museums or papers (7.4%; Fig 5c). The second most often used method of collection consisted of count surveys (34.3%). In contrast to opportunistic data collection, count surveys differ in the structure of the methodology used, and may involve transect counts, point counts, or censuses. Among other methods of collection, CS with trained volunteers only comprised a small proportion (11%) of the papers we reviewed, highlighting that training volunteers is not a barrier to publish. Nevertheless, projects with trained volunteers are more likely to be published than projects without training [52]. Data collection based on local knowledge (7.7%) and community-based monitoring (CBM) (2%) were rarely used (Fig 5c). Including local and traditional knowledge of indigenous communities in SDMs may increase the value of the data collected, the number of taxa covered and the level of engagement by participants [75, 76]. However, to fully benefit from this data collection method, researchers must be familiar with social science methods. Researchers may encounter difficulties in cross-cultural interactions, including language communication barriers and the reticence of the communities to share information about their environment (93). Despite such difficulties, both scientist and communities can benefit from building on the interest and concerns of local community members when applying local knowledge and CBM (82). Usually, the full potential of CBM programs is expressed when local communities participate actively during the entire scientific program, from the conceptual design and interpretation of results to the formulation of conclusions [75, 77]. Such cases have rarely occurred in the last decade for SDMs, probably because these programs are typically designed to monitor environmental factors rather than to collect species occurrence.

### Data type

CS data usually include presence-only (PO) data, which are easy to collect with minimum effort. In our literature review, 118 out of 224 papers used PO data, 41 used presence-absence (PA) data, and 41 used abundance (AB) data (Fig 5d). Twenty-four papers used two types of data (see S1 Table) and one paper used all three data types [78]. Several authors have highlighted the limitations of PO data, which confound information about habitat preferences and availability, have a strong spatial bias (more effort in sampling certain areas than others), and ignore environmental conditions associated with species occurrence [20, 26, 79]. If the probability of presence cannot be calculated, as in PO data (but see [80, 81]), the questions that can be addressed become limited and predictions are hindered [20, 82]. This issue with PO data explains the unexpectedly high proportion of PA and AB in the SDM papers we reviewed (47% in total; Fig 5d). Presence-absence (PA) allows the comparison of a species’ occupancy between different areas or time periods [20], but PA is generally less common in CS data. The recent development of statistical techniques to estimate occupancy, especially those that account for imperfect detection, has contributed to the increasing use of PA data to infer the spatial distribution of species [79, 83, 84]. Hence, this development would explain the increasing use of PA data obtained from CS databases. Abundance (AB) data occurred in similar proportions to PA in the set of studies reviewed. Information on the number of individuals (AB) is essential to detect changes in population sizes [20]. Presence-absence (PA) and AB data can both be obtained from checklists, point-counts, or transect surveys by volunteers [79]. The large number of studies that use PA and AB data is also highlighted by the high proportions of papers that used count-surveys data (34%, Fig 5c), and of papers that used occupancy models, generalized linear models or Bayesian hierarchical models (37.5% of the total; Fig 5e).

### Statistical approach

The statistical approaches used in the papers we reviewed were diverse, including linear regression approaches (LR, 12.7%), maximum entropy (MaxEnt, 13.3%), generalized linear models (GLM, 18.3%), occupancy models (11.2%), and generalized additive models (GAM; 10.1%; Fig 5e). Presence-only (PO) data were most frequently analyzed with MaxEnt (n = 42), whereas PA data were most frequently analyzed with occupancy models and GLM (n = 14 and 13, respectively). Abundance (AB) data were most often analyzed with GLM (n = 14). The proportion of use of the statistical approaches in CS papers that we reviewed did not seem to change over time between 2010 and 2019, with the exception of Bayesian hierarchical models (BHM) and GAMs appearing in papers from 2013 onward (Fig 5f).

### Multiple data sources

Of the 224 articles reviewed, 88 (39.7%) used multiple sources of data, merging CS with professional data. In 29 of these cases, volunteers and professional data were compared and only three studies revealed mismatches, particularly for species abundance [43, 85, 86]. The integration of CS and professional data has been a growing trend in recent years [87-89] and shows promise to improve inferences and the predictive ability of models, as well as to fill knowledge gaps for under-studied areas or poorly studied species [87-90]. This approach of combining data benefits from robust survey schemes and expands the geographic and taxonomic coverage using complementary unstructured opportunistic schemes. However, integrating highly heterogeneous data types such as large unstructured presence-only data and standardized abundance surveys, is still challenging for modelling purposes. Similar concerns exist when contemporary data are combined with historical datasets obtained from museums, grey literature or paleontological information to extend the temporal scale of a given study.

## Conclusions

In this review, we examined the trends and information gaps in the use of citizen science (CS) data for species distribution models (SDMs) in peer-reviewed papers over the last decade. Citizen science already makes substantial contributions to the field of SDMs, particularly for online occurrence databases. Indeed, the use of CS in SDMs increased exponentially during the last decade. However, taxonomic and geographic unevenness of CS projects for SDMs still remain [33, 40, 91]. This geographical disparity of data-sharing networks reduces the ability of researchers to assess national and international trends, particularly for mobile organisms [92].

The reviewed citizen science papers considered a wide range of taxa, regions, and countries, from numerous biomes and landscape types. This variability is mostly driven by the interest of the volunteers who collect data, which results in over and under-represented groups and regions. Volunteers favor certain charismatic taxa or habitats [93]. Thus, the challenge is to increase information for lesser-known locations and taxa. Filling this information gap may require reducing structured sampling efforts in areas already well-covered by volunteers, and enhancing the use of local knowledge or of community-based monitoring approaches.

We presented examples of the use of CS and highlighted recommendations to motivate further research, such as combining multiple data sources and promoting local and traditional knowledge. Accounting for the disparities in CS is crucial to adequately cover spatial and temporal scales, and strategically deploying formal surveys in areas or for species not covered by volunteers can be a key to better predict species distribution. Finally, researchers should not dismiss the impact they might have by contributing to citizen science projects. We strongly suggest researchers consider contributing to citizen science. The active participation of researchers in citizen science platforms (e.g. validating species identifications in iNaturalist) can increase the interest of participants in countries where we currently have little information on the distribution of certain species.

## Acknowledgements

This project was made possible with funding provided by the Natural Sciences and Engineering Research Council (NSERC) of Canada – Université du Québec en Abitibi-Témiscamingue (UQAT) industrial research chair on northern biodiversity in a mining context. Earlier versions of the manuscript benefited from comments by Alexandre Nolin.

## Supporting information

**S1 Table**. Methodologies of 224 papers published from 2010 to 17 October 2019 that used citizen science data to model species distribution resulting from the above described search protocol in the Scopus database.

## Author contributions

Paper selection and data collection was completed by M. Feldman. All authors made substantial contributions to all stages including the conception, research, analysis, writing and revision of this review.

